# Size-dependent tradeoffs in aggressive behavior towards kin

**DOI:** 10.1101/2020.10.26.350132

**Authors:** Chloe Fouilloux, Lutz Fromhage, Janne K. Valkonen, Bibiana Rojas

## Abstract

Aggression between juveniles can be unexpected, as their primary motivation is to survive until their reproductive stage. However, instances of aggression, which may escalate to cannibalism, can be vital for survival, although the factors (e.g. genetic or environmental) leading to cannibalism vary across taxa. While cannibalism can greatly accelerate individual growth, it may also reduce inclusive fitness when kin are consumed. As a solution to this problem, some cannibals demonstrate kin discrimination and preferentially attack unrelated individuals. Here, we used both experimental and modeling approaches to consider how physical traits (e.g. size in relation to opponent) and genetic relatedness mediate aggressive behavior in dyads of cannibalistic *Dendrobates tinctorius* tadpoles. We paired sibling, half-sibling, and non-sibling tadpoles of different sizes together in an arena and recorded their aggression and activity. We found that the interaction between size and relatedness predicts aggressive behavior: large non-siblings are significantly more aggressive than large siblings. Unexpectedly, although siblings tended to attack less overall, in size mismatched pairs they attacked faster than in non-sibling treatments. Ultimately, it appears that larval aggression reflects a balance between relatedness and size where individuals trade-off their own fitness with that of their relatives.

## Introduction

Cannibalism has often been understood to be a response to stressful environmental conditions (e.g. low-food, high competition) that is modulated by physical (e.g. size, condition) and genetic variables. It was not until the late-twentieth century that the behavior was critically evaluated and one-off observations and anecdotes (Jenkins and Carpenter, 1946; Merritt Hawkes, 1920) were synthesized into a review about the trends and occurrence of cannibalism (Fox, 1975). More recent investigations of cannibalism have recorded and tested its manifestation across a wide variety of clades, environmental states, and life history stages (Van Allen et al., 2017; Barkae et al., 2014; Van den Beuken et al., 2019; Cooper et al., 2015; Schulte and Mayer, 2017). Aggression, a behavior often tested as the precursor to cannibalistic behavior, can occur in resource-abundant, low-density environments (Fox, 1975; Mock et al., 1987; Summers and Symula, 2001), and the effect of commonly tested variables (i.e. starvation, competition) on eliciting cannibalistic behavior does not follow a consistent trend on either class-wide or even family-specific levels. Ultimately, for a behavior expressed in almost every clade in the animal kingdom, it remains difficult to tease apart the evolutionary motivation behind cannibalism.

Aggression in cannibals can be difficult to interpret, as the motivation is not always hunger-driven, and death is not always the end-state of aggressive interactions. For example, cannibalistic lizards (*Podarcis gaigeae*) in Greece show local variation in aggression, where males are more aggressive in high-density (but high-resource) islands. It appears that aggression by males leads to the consumption of conspecific tails, which although is not always deadly, serves as a high-fat meal that decreases the sexual quality of male competitors (Cooper et al., 2015). Aggression between juveniles complicates matters further, as individuals are not competing for mates and usually do not hold territories. For the most part, juvenile aggression and cannibalism are justified by competition for immediate nutritional resources (larval flounders: Dou et al., 2000; earwig nymphs: Dobler and Kölliker, 2011), or parental care (vulture chicks: Margalida et al., 2004) which may be survival mechanisms to compensate for fluctuating environmental conditions (Mock et al., 1987).

Theoretical justifications predicting cannibalism are often built around physical qualities. Across the animal kingdom, many studies have found that cannibals are most often larger than their prey (Barkae et al., 2014; Claessen et al., 2004; Ibáñez and Keyl, 2010; Rojas, 2014) although exceptions exist when larger individuals are weakened (Richardson et al., 2010). Kinship between individuals has also been used to characterize cannibalistic interactions. This has been shown to be an important factor in several cannibalistic species who demonstrate kin discrimination and avoid eating kin (salamanders: Pfennig et al., 1994, bulb mites: Van den Beuken et al., 2019, toad tadpoles: Pfennig and Frankino, 1997), though this is far from a rule, as similar range of cannibals do consume their kin without avoidance (moths: Boots, 2000, poison frog tadpoles: Gray et al., 2009). Despite kinship and physical attributes driving much of the discussion on the evolution and maintenance of cannibalism, few studies have incorporated both in predicting cannibalism outside of oophagy or sexually competitive contexts. Moreover, such studies have so far been limited to invertebrates: in European earwigs, kinship and weight asymmetry affected cannibalistic behaviour both separately and in an interaction, such that weight asymmetry effects were stronger among unrelated individuals (Dobler and Kölliker, 2011). In both desert (Bilde and Lubin, 2001) and wolf spiders (Roberts et al., 2003), kinship but not weight asymmetry affected cannibalism. While fascinating in their own right, these studies hardly provide a basis for generalising to distant clades such as vertebrates.

*Dendrobates tinctorius* is a species of Neotropical poison frog whose larvae are aggressive cannibals (Rojas, 2014). Tadpoles are often deposited by their fathers in ephemeral pools of water, where they are left to develop until metamorphosis (Rojas and Pašukonis, 2019). Although tadpoles are most often transported singly, the ephemeral pools in which they are deposited can have multiple tadpoles of various developmental stages (Rojas and Pašukonis, 2019) and degrees of relatedness (B. Rojas & E. Ringler, unpublished data). In these environments, cannibalism is common (Rojas, 2014, 2015), yet the escalation of aggression to cannibalism in this species has not been explicitly tested. Closely related poison frogs have linked aggressive behavior and cannibalism (Gray et al., 2009; Summers and Symula, 2001), though exceptions exist in obligate egg-feeders with parental care (Dugas et al., 2016). For *D. tinctorius*, the costs of aggression are direct, as consuming kin reduces inclusive fitness and the potential for injury (even with a small counterpart) is high; the potential benefits, on the other hand, are complex, as by shortening time to metamorphosis and increasing physical size thereafter, individuals may be able to escape precarious conditions and improve fitness prospects.

In this study we tested physical and genetic variables to better understand the basis of aggression in a cannibalistic species. We conducted behavioral assays between pairs of *Dendrobates tinctorius* tadpoles, and measured aggression and activity in response to size differences and relatedness. These tests will allow us to evaluate the importance of physical characteristics with respect to genetic relatedness as predictors of aggression. Experiments were supplemented with theoretical models based on inclusive fitness theory to study the predictors of aggression in this species; together, these experiments and models contribute to our understanding of how cannibalism is shaped by the costs and benefits of relatedness and aggression in animals.

## Methods

### Study species

*Dendrobates tinctorius* is a species of Neotropical poison frog with elaborate parental care. Males attend small terrestrial clutches and transport newly-hatched tadpoles, one or two at a time, to pools of water where they are left until metamorphosis. Males carrying more than one tadpole at once can be seen either depositing both tadpoles in the same pool or distributing tadpoles between pools (Rojas and Pašukonis, 2019). Tadpoles of this species are omnivorous and frequently demonstrate cannibalistic behaviour (Rojas, 2014, 2015); despite this, it is not unusual to see tadpoles of various stages coexisting within the same pool in the wild (Rojas and Pašukonis, 2019).

We used tadpoles from a breeding laboratory population of *D. tinctorius* kept at the University of Jyväskylä, Finland. We maintained a paternal half-sibling design as it could be expected that paternal half-siblings are more likely to co-occur as a result of fathers reusing pools after multiple transport events. Tadpole dyads were assigned in response to (1) individuals needing to be visually distinguishable from each other (i.e. size), and (2) the laboratory mating schedule/network, which was prioritized as to not stress the animals from overbreeding. Adult pairs were each housed in a 55L terrarium that contained layered expanded clay, leaf-litter, moss substrate and were equipped with a shelter, logs, and live plants. Terraria were maintained at 26C (±2C) and were automatically misted with reverse osmosis water four times a day (maintaining a humidity around 95%) and lit with a 12:12 photoperiod. Frogs were fed live *Drosophila* fruit flies coated in vitamin supplements five times per week. Tadpoles were raised singly in 10 x 6.5 x 5 cm containers which were filled with spring water, and fed *ad libitum* a diet of fish food (JBL NovoVert flakes) three times a week. Adult and tadpole health and water levels were checked daily.

### Behavioral trials

Pairs of tadpoles of different degrees of relatedness (full sibling, half-sibling, non-sibling) were placed together in an arena. Tadpoles in early larval development were used, but we established a cut-off point before the toe differentiation in hind legs development to control for possible life-history effects (stage 31; Gosner, 1960). Experimental tadpole weight ranged from 0.04g to 0.38g, and size differences between pairs ranged from 0.03g to 0.30g. Blinding in the experiment was not possible, as the set-up and experiment were conducted by the same person, but the order of trials was assigned randomly. The arena was a 18.5cm by 12cm clear plastic container filled with 400 mL of spring water; four quadrants were delineated on the base of the arena to provide information about tadpole activity throughout the experiment. Initially, each tadpole was placed on either side of an opaque partition dividing the arena; this partition kept tadpoles separated but allowed water to flow throughout the container. After an acclimation period of one hour, tadpole activity (resting, swimming) of the separated individuals was recorded every 15 seconds for 10 minutes.

Following the acclimation and separated observation, the barrier was removed and tadpole interactions were recorded for 60 minutes. Focal behaviors (resting, swimming, biting, and chasing; see Table 1 for descriptions) were recorded for both tadpoles every 15 seconds. Tadpoles were visually distinguishable from each other as a result of size differences. Individuals were photographed and weighed before the beginning of each trial to establish initial tadpole condition, and were only used once (n_Trial_ = 15 for each relatedness level, n = 90 tadpoles for the entire experiment).

**Table 1.**
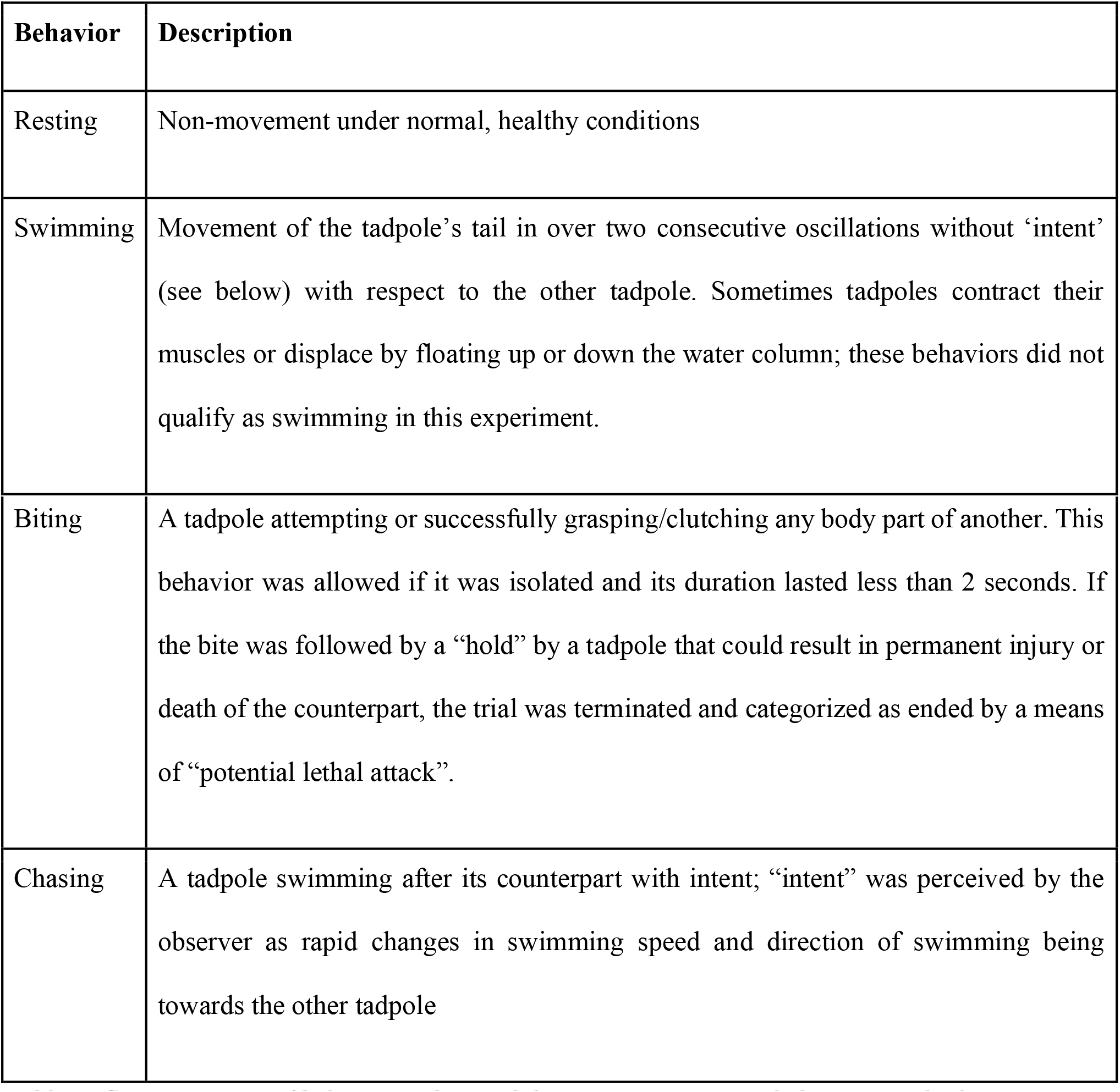
Categorization of behaviour observed during aggression trials between tadpoles.

| Behavior | Description |
| --- | --- |
| Resting | Non-movement under normal, healthy conditions |
| Swimming | Movement of the tadpole’s tail in over two consecutive oscillations without ‘intent’ (see below) with respect to the other tadpole. Sometimes tadpoles contract their muscles or displace by floating up or down the water column; these behaviors did not qualify as swimming in this experiment. |
| Biting | A tadpole attempting or successfully grasping/clutching any body part of another. This behavior was allowed if it was isolated and its duration lasted less than 2 seconds. If the bite was followed by a “hold” by a tadpole that could result in permanent injury or death of the counterpart, the trial was terminated and categorized as ended by a means of “potential lethal attack”. |
| Chasing | A tadpole swimming after its counterpart with intent; “intent” was perceived by the observer as rapid changes in swimming speed and direction of swimming being towards the other tadpole |

Trials were ended prematurely if tadpoles demonstrated aggression levels that would cause severe damage or death (where bites lasted for more than 2 seconds, recorded as “potential lethal attack”). Although aggression was common, potential lethal attacks were rare, occurring in only 3/45 trials. There were no tadpole deaths as a result of the behavioral trials, and all tadpoles were kept and reared in the laboratory after the experiment. Assays were done according to the Association for the Study of Animal Behaviour’s guidelines for the treatment of animals in behavioural research and teaching (ASAB 2017), and with the approval of the National Animal Experiment Board at the Regional State Administrative Agency for Southern Finland (ESAVI/9114/04.10.07/2014).

**Image 1.**
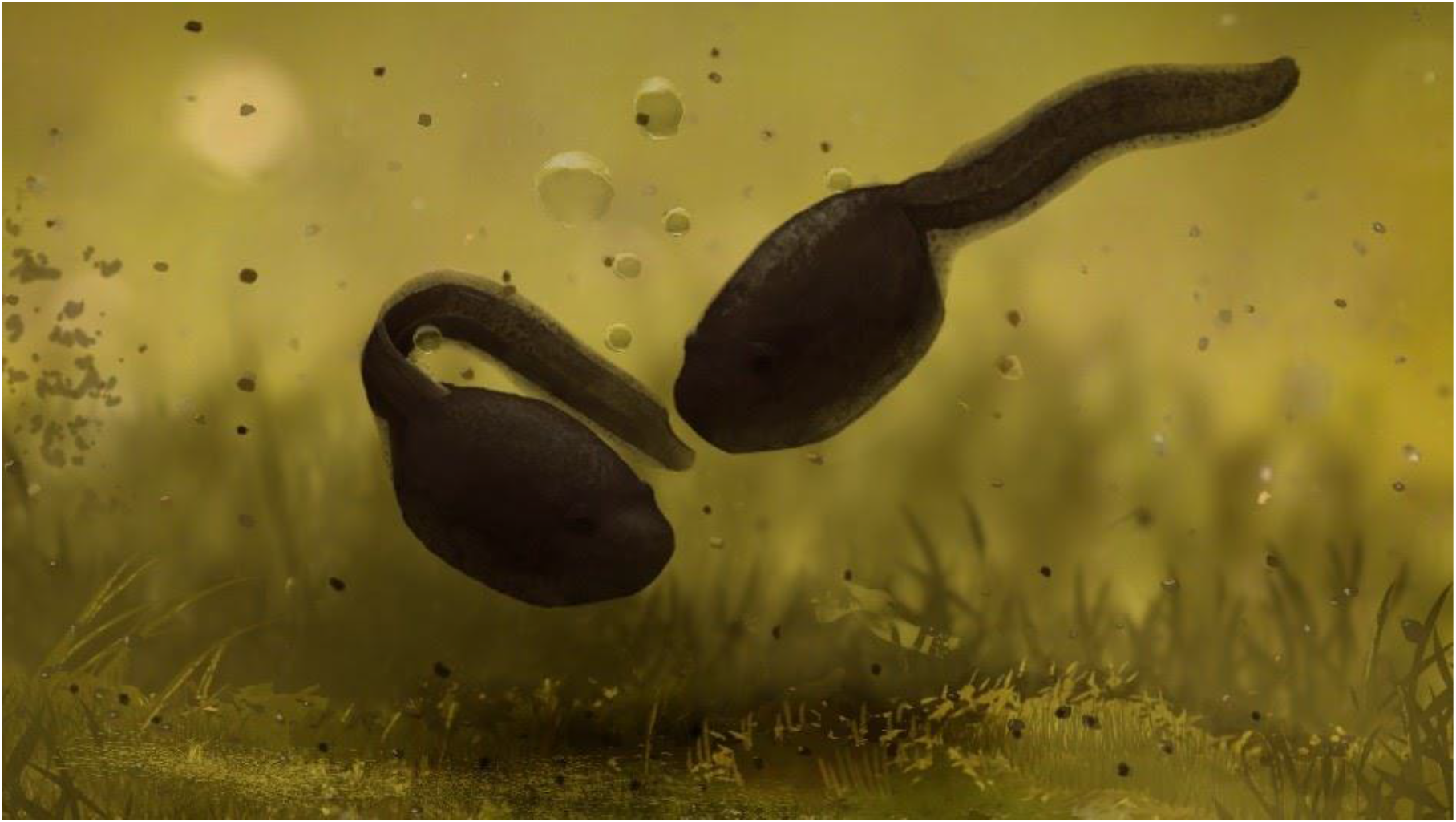
Illustration of aggressive exchange between D. tinctorius tadpoles. Note the tail bite on one of the tadpoles, which is a common occurrence both in wild and laboratory encounters. Published with artist permission.

### Statistical analysis

All models and statistics were performed in the program R (v. 3.6.1, R Development Core Team, 2019) with additional packages “glmmTMB” (Magnusson et al., 2020), “coxme” (Therneau, 2020), “dplyr” (Wickham et al., 2018), “tidyr” (Wickham et al., 2019). Activity and aggression analyses (see below) took into account pair identity (Pair_ID) and family (Breeding pair) level random effects (Supp. tables 1 and 2 for AIC comparisons). Differences in duration of trials during experiments (n = 3/45 trials ended early due to potential lethal attacks) were taken into account by offsetting models with a trial duration. Aggression and activity models were chosen based on the Akaike Information Criterion (AIC, Akaike, 1973), which compares fit based on log-likelihood and the number of model parameters (see supplementary materials for model AIC comparisons). We considered both negative binomial (with linear and quadratic parameterization) and poisson distributions as model families in AIC comparisons. Residual diagnostics and overdispersion were checked and corrected using the “DHARMa” (Hartig, 2020) package.

### Activity levels

Tadpole activity was categorized as “resting” and “swimming” (see Table 1 for details). Tadpole activity was observed during post-acclimation (10 minutes) and experimental (max. 60 minutes) periods. Activity was coded as counts and were modeled in a generalized linear mixed model framework (GLMM). The best fitting model used a negative binomial linear parameterization where activity was predicted by relative tadpole size (a categorical variable (i.e. small, large), where size is relative within pairs) and relatedness (a categorical variable with three levels: full sib, half sib, non sib) (see Supplementary Table 1).

**Table 1.** Summary of negative binomial GLMM with linear parameterization of tadpole activity chosen from AIC model comparison. Where **(A)** half-siblings demonstrated significantly less activity compared to siblings and non-siblings while tadpoles were separated during post-acclimation, which changes during the experiment **(B)** where large tadpoles are overall more active. Models for (A) and (B) were predicted by additive effects of size and relatedness. Tadpole dyads (Pair_ID) and family were accounted for as random effects, CI represents 95% confidence interval. Differences in trial time during the experiment (n = 3/45) were accounted for by using duration as offset in the model. Experimental activity model was overdispersed and size was corrected for, hence missing residual variance (σ^2^) in Panel B; τ_00_ represents random intercept variance.

| **A.** |  |  |  |  | **B.** |  |  |  |  |
| --- | --- | --- | --- | --- | --- | --- | --- | --- | --- |
| **Post-acclimation Activity** |  |  |  |  | **Experimental Activity** |  |  |  |  |
| *Predictors* | *Estimate* | *CI* | *z* | *p* | *Predictors* | *Estimate* | *CI* | *z* | *p* |
| (Intercept) | 0.99 | 0.30 – 1.67 | 2.82 | **0.005** | (Intercept) | 0.00 | 0.00 – 0.01 | -29.09 | **<0.001** |
| Halfsiblings | -0.94 | -1.82 – -0.06 | -2.08 | **0.037** | Halfsiblings | 1.08 | 0.76 – 1.53 | 0.43 | 0.665 |
| Siblings | -0.19 | -0.94 – 0.55 | -0.50 | 0.616 | Siblings | 0.97 | 0.68 – 1.39 | -0.16 | 0.874 |
| Size (Large) | -0.07 | -0.69 – 0.55 | -0.22 | 0.824 | Size (Large) | 2.53 | 1.79 – 3.58 | 5.23 | **<0.001** |
| **Random Effects** |  |  |  |  | **Random Effects** |  |  |  |  |
| $\sigma^2$ | 1.71 | | | | $\sigma^2$ | | | | |
| $\tau_{00}$ Pair_ID | 0.14 | | | | $\tau_{00}$ Pair_ID | $8.43 \times 10^{-9}$ | | | |
| $\tau_{00}$ Family | $1.82 \times 10^{-8}$ | | | | $\tau_{00}$ Family | $5.76 \times 10^{-3}$ | | | |

### Overall aggression

Aggression between tadpoles was observed as chasing or biting (see Table 1), which were recorded as counts. These two behaviors were combined to represent “total aggression”. The best fitting model for aggression was a generalized linear mixed effect model (GLMM) with a negative binomial linear parameterization. Total aggression was predicted by relatedness (a categorical variable with three levels: full sib, half sib, non sib) and size (a categorical variable (i.e. small, large), where size is relative within pairs)(see Supplementary Table 2), pairs were taken into account as a random variable. We had modeled mass difference (a continuous predictor) between tadpoles as potential predictors which resulted in less parsimonious models (see Supplementary Table 2).

### Latency to first bite

We modeled latency to first biting behavior using a mixed effect Cox proportional hazards model. Survival object was parameterized with respect to latency to first bite event and absolute biting (0/1, where 0 represents no biting occurred during the trial) in response to the interaction of relatedness and mass difference between tadpole dyads. Latency data was built by selecting the “first biter” within a pair which involved subsetting the original data set. As a result of this, we used mass difference between large and small tadpoles (instead of categorical size) to incorporate trials without aggressive behavior. Mass difference was calculated as the difference between the large and small tadpoles within pairs: this value was always positive since large tadpoles were always more massive. Using subsetted data, each pair identity was independent, so only “Family” was used as a random variable.

### Game theory model

We modeled pairwise interactions between tadpoles arbitrarily labelled as 1 and 2. We assumed that only one tadpole per pair survives (‘wins’), and that the probability of winning depends on each individual’s competitive strength. Competitive strength *θ_i_* of tadpole *i* was calculated based on its **size**, *s_i_* and its **aggressiveness,** *a_i_* as *θ_i_* = *s_i_* · *a_i_*. This multiplicative formulation reflects the biological idea that a given increment in aggressiveness should have a greater effect on a large than a small tadpole’s competitive strength. Individual 1’s probability of winning is given by its **relative competitive strength**, as 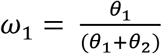. The reproductive success (“**direct fitness**”, *ν_i_*) of the winning tadpole was modeled under three alternative assumptions: (*1a*) *ν_i_* is size-independent, as *ν_i_* = 1 – *a_i_*^2^; (*1b*) *v_i_* is proportional to size (for a given level of aggressiveness), as *ν_i_* = *s_i_* – *a_i_*^2^; and (*1c*) *ν_i_* is sizedependent due to aggressiveness being costlier for smaller tadpoles, as 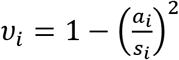 (see Fig 3 for visualization). In all three formulations costs increased at an accelerating rate, such that low levels of aggression had low costs whereas high levels of aggression could be extremely costly; this was done to account for the increasing danger and energy expense associated with more violent behaviors.

**Figure 1.**
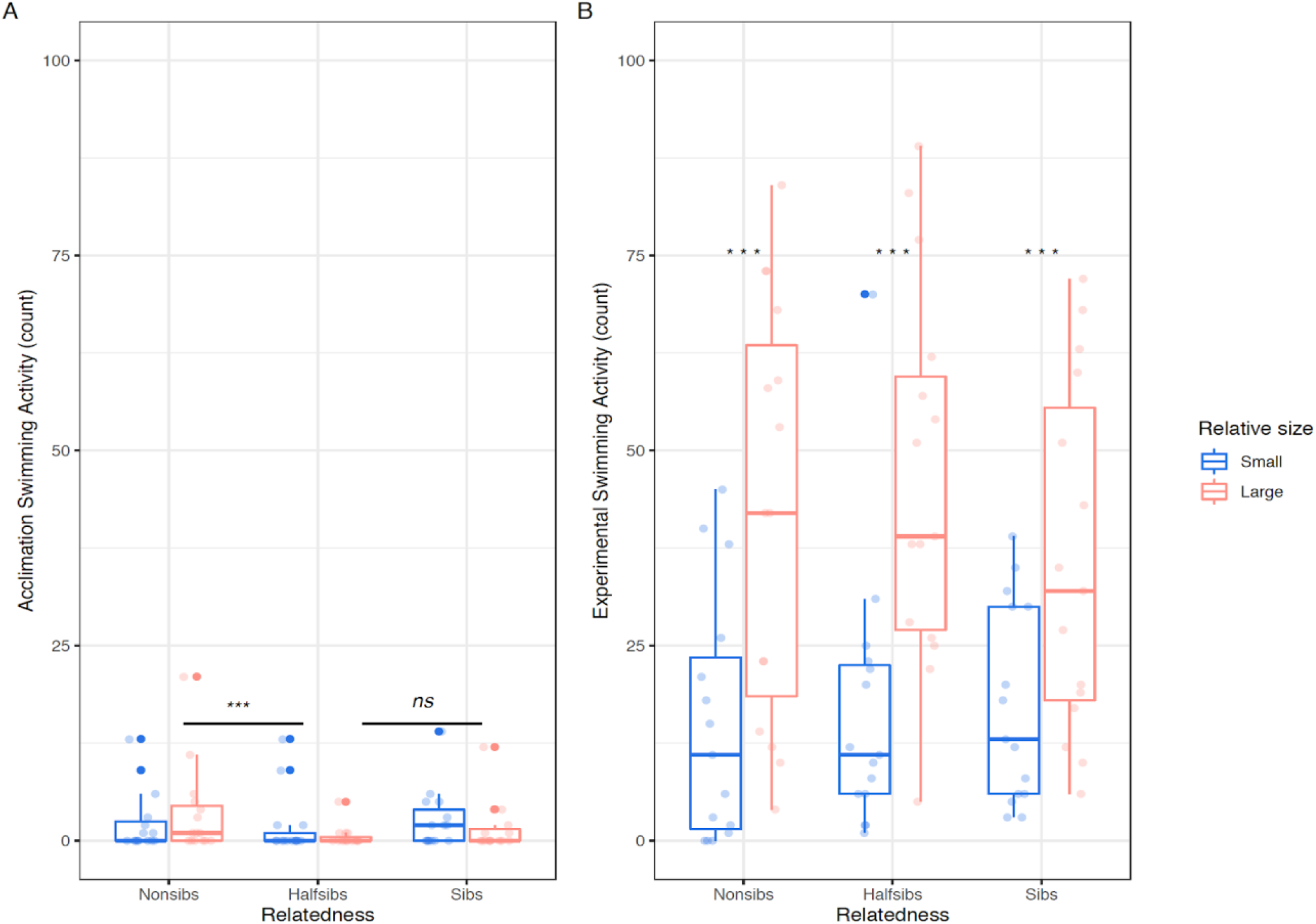
Tadpole activity levels before and during experimental trials. Panel (**A**) shows the post-acclimation activity of tadpoles. We found no difference in swimming activity between large and small tadpoles during this phase, but found that half siblings were significantly less active during this period. Panel (**B**) shows experimental activity throughout behavioral trials. Large tadpoles were significantly more active than small tadpoles during assays. N_Trial_ = 15 for each relatedness level. Large tadpoles are in pink and small tadpoles in blue. Boxplot medians are depicted by thicker lines, whiskers span ± 1.5 * interquartile range.

**Figure 2.**
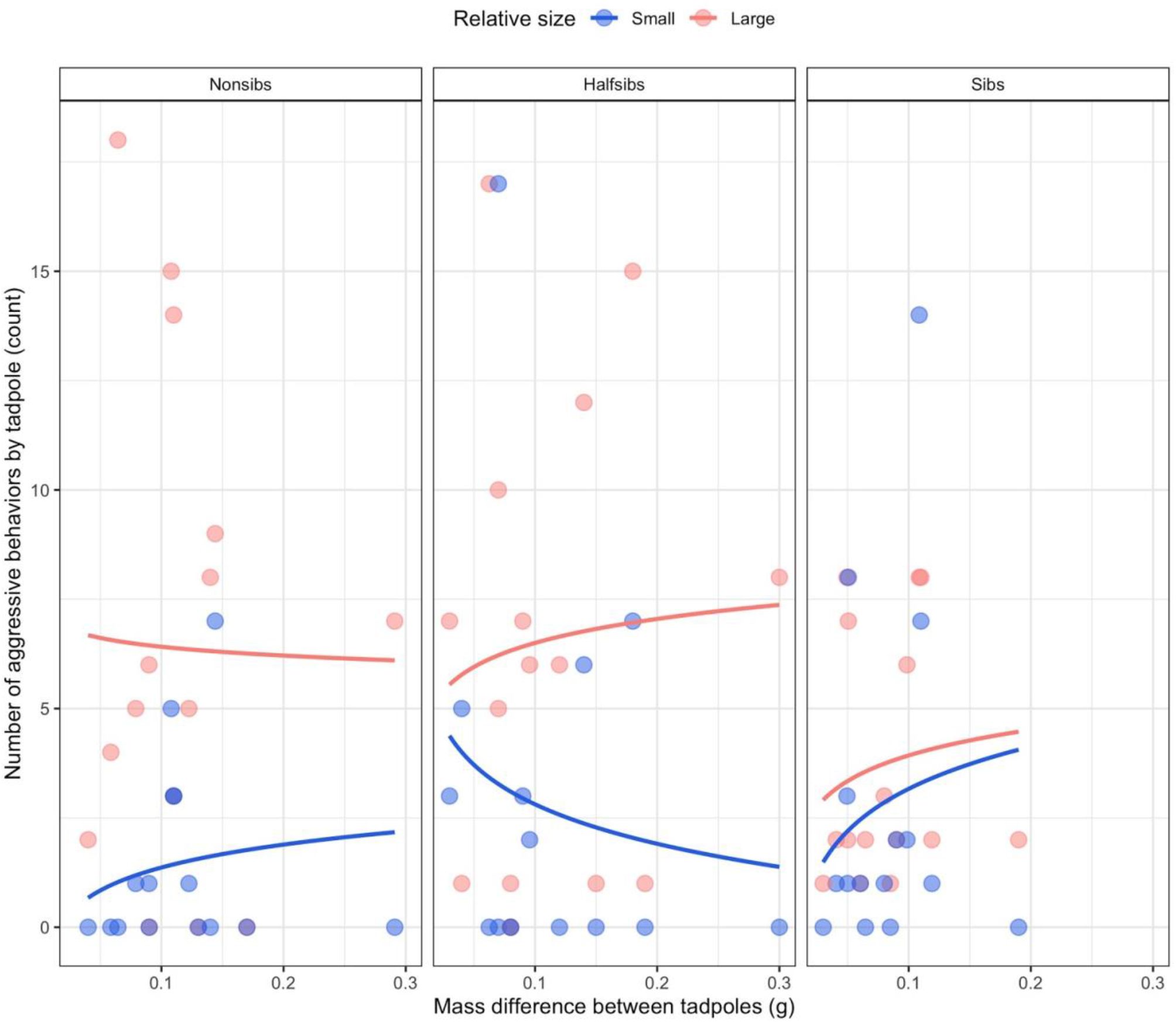
Differences in aggression across relatedness treatments with respect to weight difference between dyads. Pink dots represent large tadpoles and blue dots represent small tadpoles (sizes relative to dyads); N_Trial_ = 15 for each relatedness level. There was a higher level of aggression by large tadpoles overall, but significantly less aggression by large siblings when compared to non-siblings. Lines fit with GLM smoother (y ~ log(x)) for relative sizes.

**Figure 3.**
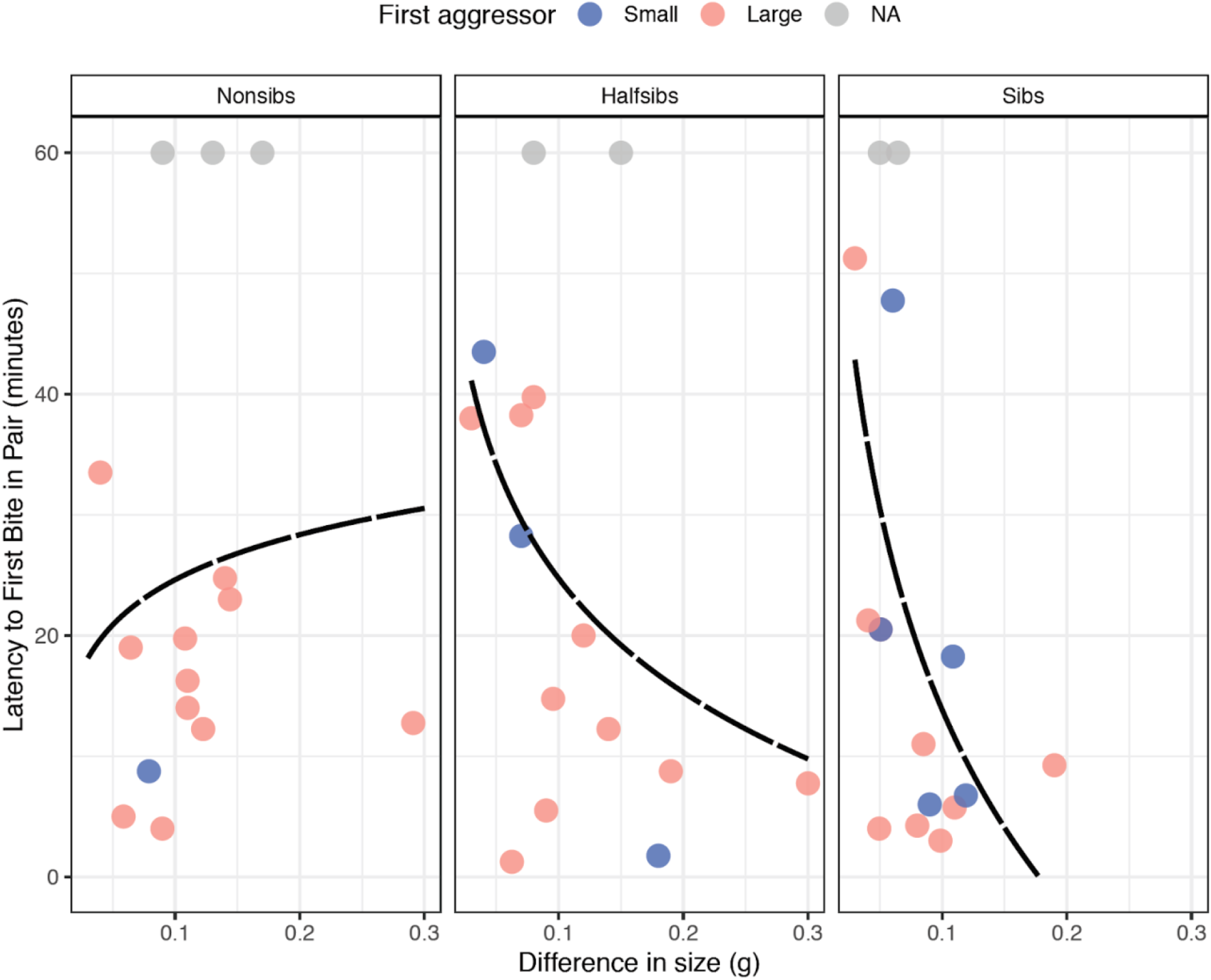
Latency to first bite between tadpole dyads. Points are colored by relative tadpole size within dyads. Lines are fit with a GLM smoother with a y ~ log(x) formula. There is an inversion in behavior as size difference increases, where sibling pairs with large size differences attacked significantly faster than non-siblings. Dyads where there were no aggressive behaviors were accounted for by assigning them the maximum time limit (60 minutes). N_Trial_ = 15 for each relatedness level.

Finally, the inclusive fitness of the surviving tadpole was calculated as *ν*_1_ – *r ν*_2_, where *r* is the relatedness between the pair. This formulation reflects the idea that winning involves the killing of a relative who would have had reproductive success ν_2_ had it survived. The inclusive fitness of the losing tadpole is zero, because the losing tadpole neither reproduces nor affects the other tadpole’s reproduction. We calculated the expected (i.e., probability-weighted mean) inclusive fitness of tadpole 1 as *F*_1_ = *ω*_1_(*ν*_1_ – *r ν*_2_). For given values of *s*_1_, *s*_2_, and *a*_2_ we numerically determined individual 1’s optimal aggression level as the value of α_1_ that maximises its expected inclusive fitness. By computing individual 1’s ‘best response’ aggression level for any given *a_2_* which its opponent might exhibit, we then identified pairwise optimal aggression levels that are best responses to each other.

## Results

### Activity levels

We observed tadpole activity during both post-acclimation and experimental phases. While tadpoles were separated by an opaque barrier during the post-acclimation phase (but water still freely moved throughout the arena) we found that half-siblings demonstrated significantly less swimming behavior than non-siblings (negative binomial GLMM, CI: 0.06-1.82, *z* = −2.08, *p* = 0.037) but not siblings (negative binomial GLMM, CI: −0.15−1.65, *z* = 1.64, *p* = 0.102). During the experiment, however, we found that large tadpoles across all relatedness treatments were significantly more active than small tadpoles (negative binomial GLMM, CI: 1.79 - 3.58, *z* = 5.23, *p* < 0.001; see Fig 1, Table 1).

When comparing models we found that random effects of pair ID had higher between subject variance (τ_00_ = 0.14) than tadpole family (τ_00_ = 1.82 x 10^-8^) during post-acclimation activity (Table 1, Panel A), indicating that when separated, there was less variation in behavior on a family level. Yet, while interacting during the experiment this difference disappears (Table 1, Panel B). In both cases, between-subject variance is low, indicating that across families and pairs of tadpoles, activity levels are similar.

### Overall aggression

We found that large siblings were significantly less aggressive than the large non-sibling treatment, exhibiting almost half the amount of aggressive behaviors as large non-siblings (Fig. 2, negative binomial GLMM, *z* = −2.07, *p* = 0.039, Table 2). Half-siblings were not significantly different from either treatment. Following our expectations of creating unique pair interactions, the random effect of pair identity had a high between group variation (τ_00Pair_ID_ = 0.55, Table 2), but families were unexpectedly consistent (τ_00Family_ = 0.06, Table 2).

**Table 2.** Summary of negative binomial GLMM with linear parameterization of tadpole aggression. Where total aggression (total count of biting and chasing) was predicted by the interaction between size (two level categorical variable) and relatedness. Tadpole dyads (Pair_ID) and family were accounted for as random effects, CI represents 95% confidence interval. Differences in trial time during the experiment (n = 3/45) were accounted for by using duration as offset in the model. σ^2^ represents residual variance and τ_00_ represents random intercept variance.

| *Predictors* | **Total aggression** |  |  |  |
| --- | --- | --- | --- | --- |
|  | *Estimate* | *CI* | *z* | *p* |
| (Intercept) | -7.95 | -8.75 – -7.16 | -19.63 | **<0.001** |
| Halfsiblings | 0.41 | -0.62 – 1.44 | 0.77 | 0.440 |
| Siblings | 0.58 | -0.40 – 1.56 | 1.15 | 0.249 |
| Size (Large) | 1.44 | 0.68 – 2.20 | 3.71 | **<0.001** |
| Halfsiblings: Size (Large) | -0.44 | -1.47 – 0.58 | -0.84 | 0.399 |
| Siblings: Size (Large) | -1.02 | -1.99 – -0.05 | -2.07 | **0.039** |
| **Random Effects** |  |  |  |  |
| $\sigma^2$ | 0.58 | | | |
| $\tau_{00}$ Pair_ID | 0.55 | | | |
| $\tau_{00}$ Family | 0.06 | | | |

### Latency to first bite

We observed that initial aggression between tadpoles can change based on physical and genetic attributes. We used biting behavior as a measurement of first aggression because it consistently represented the first aggressive contact in tadpole dyads. Based on a mixed effect Cox proportional hazards model, we assessed the risk of first attack when considering relatedness and size difference between pairs. We found a significant interaction between relatedness and size, where a higher degree of relatedness led to a shorter latency to aggression when the size difference within dyads was larger. In other words, siblings demonstrated more immediate aggressive behavior towards each other when their size differences were greater (Cox mixed effects, *z* = 2.209, *p* = 0.022, see Table 3). For example, at a large mass difference (> 0.15 g between tadpoles) siblings were more than 40 percent more likely to bite than non-siblings within the first five minutes of a trial. Interestingly, non-siblings demonstrated a seemingly inverted behavioral trend, where dyads with large size differences had delayed aggressive behaviors. Half-siblings did not behave significantly differently from either treatment. In trials where biting was exhibited, large tadpoles were most often the first aggressor (n = 8/13 for siblings; n= 10/13 for half siblings; n= 11/12 for non siblings).

**Table 3.**
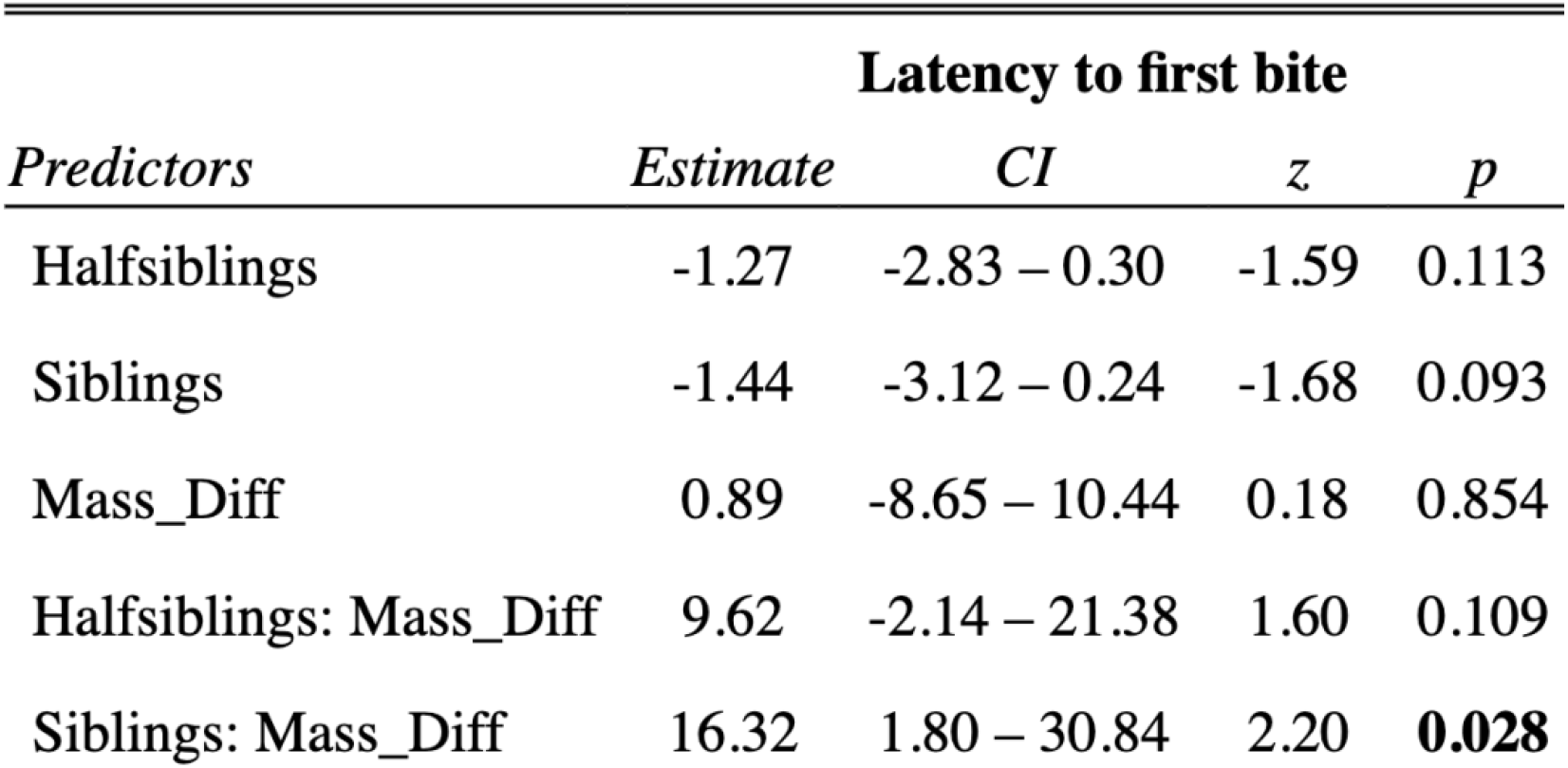
Mixed effects Cox proportional hazards model. Time to first aggressive behavior was predicted by the interaction of the mass difference between tadpoles and their relatedness; family is taken into account as a random effect. There is a significant interaction between relatedness and size, where more similarly sized siblings have a shorter latency to aggression than non-siblings. Mass_Diff is the difference in weight between large and small tadpoles.

### Game theory model

Based on our three formulations (1a-c) we varied the impact of size to model aggression levels of tadpoles with different degrees of relatedness. The version where aggression was both size-dependent and costlier for the smaller tadpoles (Fig 4, third row) appeared most consistent with our empirical data (Fig 2), in that larger tadpoles were consistently predicted to be more aggressive than their smaller counterparts, and overall aggression decreased with relatedness.

**Figure 4.**
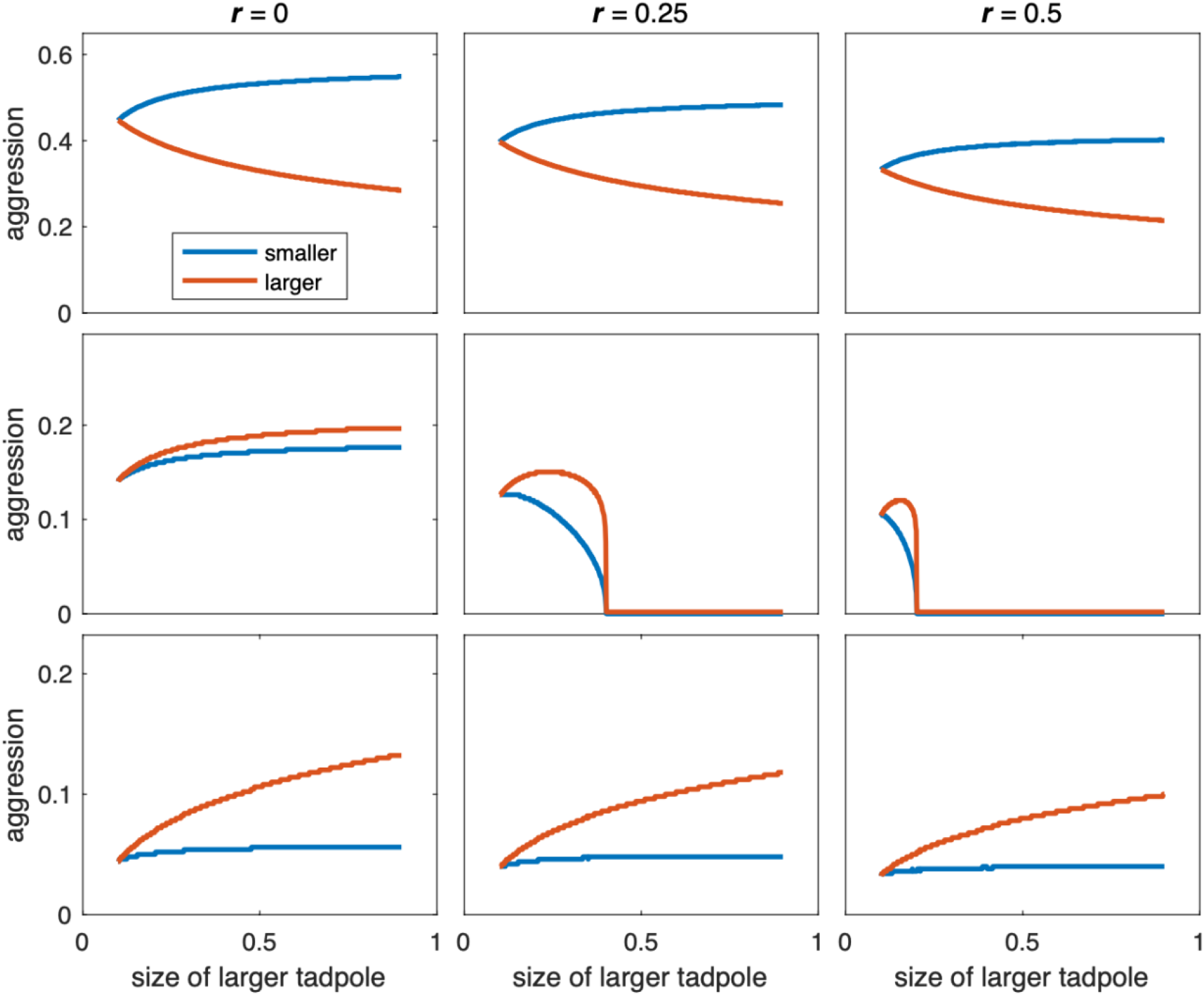
Optimal aggressiveness of dyads of tadpoles as a function of size difference for three different levels of relatedness (represented in panel columns) and three sets of assumptions (represented in panel rows). **First row:** direct fitness was assumed to be size-independent. **Second row**: direct fitness was assumed to be proportional to size. **Third row**: aggressiveness was assumed to be costlier for smaller tadpoles. The smaller tadpole’s size was held fixed at si = 0.1; the larger tadpole’s size is shown on the x-axis.

## Discussion

Larval cannibalism or, more broadly, juvenile aggression has most often been studied in the context of resource competition (Baras et al., 2000; Tershy et al., 2000), as juveniles do not usually defend territories (Stamps and Krishnan, 1994), nor compete for sexual partners. Here, we observed aggressive behaviors between *D. tinctorius* tadpoles in resource-abundant, low-density conditions. We found that aggression is common (also see Fischer et al., 2020; Rojas, 2014), and depends on the interaction between relative size and relatedness between tadpoles. Our results show that, together, relatedness and physical attributes shape overall aggression, latency to aggression, and even activity levels between dyads.

### Cannibalism and the environment

Aggressive attacks between pairs were recorded across all relatedness treatments and sizes. Although less common, small tadpoles were sometimes quicker to exhibit aggressive behaviors than their larger counterparts (Fig 3) and, in some instances, were even more aggressive than large tadpoles (this was observed only in sibling and half-sibling treatments, see Supplementary Fig 1). These intense aggressive responses were elicited under vastly different conditions from which cannibalism is usually reported, such as in response to starvation (spiders: Mayntz and Toft, 2006; cephalopods: Ibáñez and Keyl, 2010; earwigs: Dobler and Kölliker, 2011), pathogens (caterpillars: Wang and Daane, 2014; salamanders: Pfennig et al., 1991), high population densities (crabs: Moksnes, 2004), and in the context of sexual selection (chimpanzees: Takahata, 1985; seals: Bishop et al., 2016). These associations suggest that cannibalism is triggered by immediate stress and competition. However, a handful of studies have been invoked in decoupling stress from aggression and cannibalism (poison frogs: Dugas et al., 2016b; egrets: Mock et al., 1987, vultures: Margalida et al., 2004, snails: Baur, 1987), contributing to the mounting evidence that across diverse taxa, immediate physical and environmental stress are not the only triggers for cannibalistic behavior (Fox, 1975).

We report here that aggression in *D. tinctorius* is not primarily driven by immediate nutritional need. This raises the question of whether there is a long-term fitness advantage to a cannibal that pays off later, during the reproductive stages of the animal. For example, attackers may accrue fitness advantages by having a shorter latency to metamorphosis, larger size as an adult, and higher fecundity as an outcome of their behavior during the larval stage (Crump, 1990; Polis, 1981).

### Interactive predictors of cannibalism

Aggression between tadpoles was predicted by the interaction between relative size and relatedness within dyads. We found that large non-siblings were the most aggressive, expressing almost twice the amount of aggressive behaviors than large siblings (Fig. 2). When able to physically interact in the arena, the behavior of half-siblings did not differ significantly from the two other relatedness treatments, which demonstrates that even if there does appear to be some kin discrimination in *D. tinctorius*, it may not function on as fine of a scale as for other species (e.g., Pfennig et al., 1994).

With respect to the interaction with size, Rojas (2014) had already established that cannibalism between *D. tinctorius* tadpoles occurs faster with increasingly size-mismatched pairs. In fact, across the animal kingdom the aggressor in a pair/group is most often the larger individual (Ibáñez and Keyl, 2010; Mayntz and Toft, 2006; Mock et al., 1987). However, our findings highlight the fact that in this system aggression is not solely mediated by size differences. This suggests that there may be many other cases of context-specific discrimination across taxa that have been overlooked by not considering the interaction between relatedness and physical attributes (such as illness, injury, or phenotype). The context-specific kin discrimination observed in this study could have potentially evolved because the benefit to cannibals only outweighs the cost of harming kin in particular circumstances.

In the salamander system, the cost of consuming kin is high (i.e. disease rates, Pfennig, 1999; Pfennig et al., 1991), thus the ability to discriminate against them is valuable; even first cousins have been shown to be attacked less than nonrelatives (Pfennig et al., 1994). Costs may be lower for *D. tinctorius* tadpoles if they do not face similar consequences of cannibalism such as acquiring pathogens (although the effects of pathogens on cannibalism in this species remain unexplored). Costs may also be decreased by the impact of parental care, whose protective effects have been shown in other poison frog species (Schulte et al., 2011; but see Rojas, 2014, 2015) for seemingly counterintuitive deposition choices in *D. tinctorius*). Although discrimination is less precise than in salamanders (Pfennig et al., 1994), our results support the presence of kin discrimination in *D. tinctorius*, and warrant further investigation into the possible proximate mechanisms regulating kin recognition in this species.

### Latency to aggressive behavior

Latency to attack changed unexpectedly as a function of both size difference and relatedness between tadpoles. When pairs were more similarly sized, non-siblings attacked faster; in contrast, when mismatched in size, non-siblings delayed aggression (Fig. 3). This trend was inverted for siblings, which were tolerant of a similarly-sized counterpart, but were quickly aggressive in size-mismatched pairings. In other words, although large siblings were less aggressive overall, they had a shorter latency to aggression in dyads with large size differences. We speculate that fast ‘attacking’ may serve different functions in different contexts. For example, when performed between size-mismatched siblings, it may serve not to initiate an escalated fight but to ascertain by taste the first impression of relatedness.

When considering latency to aggression, it may be important to take into account at which point tadpoles processed that they were not alone in the arena. The earliest occasion where this could have occurred is during the acclimation phase, during which tadpoles were separated by an opaque barrier which allowed water to pass through the entire testing arena (for an hour during acclimation, and actively recorded for 10 minutes before the experiment). It is probable that chemical cues (if any) and vibrations from tadpoles moving in the water circulated throughout the entire arena during this time. Unexpectedly, half siblings were significantly less active than non-siblings and did not differ in activity from siblings while dyads were separated (Fig 1, Panel A); relatedness-level differences in activity disappeared once pairs could physically interact (Fig 1, Panel B). It should be noted that post-acclimation behaviour was observed for only 10 minutes, and that tadpoles could have behaved differently during the unobserved acclimation period (60 minutes). Nevertheless, higher activity levels by non-siblings could indicate the presence of some kind of chemical cue whose meaning then shifted with the presence of visual contact.

While we are unsure of the mechanisms *D. tinctorius* may be using to recognize each other, we believe that tadpoles could potentially be using both olfactory and taste cues to discriminate kin, as shown in salamanders (Pfennig et al., 1994) and *Xenopus* sp. (Dulcis et al., 2017). Fischer et al., (2020) recently described the neural basis of conspecific aggression in *D. tinctorius* and found differences in brain region activity based on the “winning” or “losing” status of paired tadpoles after fights; their findings lay the groundwork for understanding the proximate mechanisms of aggression and provide the backdrop to understanding its triggers in *D. tinctorius* tadpoles. Following the establishment of kin discrimination in the species (this study), experiments investigating the possible mechanisms underlying recognition are warranted.

### Game theory model

As intuitively expected, our model robustly predicts decreasing aggression levels with increasing relatedness (Figure 4). Predictions about size-dependent behavior, however, turn out to be sensitive to details. When aggression is assumed to be costlier for small tadpoles (bottom row of panels in Fig. 4), larger tadpoles are always more aggressive, and are increasingly so towards larger size differences and more distant relatedness. This formulation (1c) results in a pattern that is the most consistent with experimental results (Fig 4). The model also reveals interesting theoretical possibilities which can be rejected for *D. tinctorius*. For example, if (adult) reproductive success and aggressiveness costs were independent of tadpole size, smaller tadpoles should compensate for their size disadvantage by being more aggressive (top row of panels in Fig. 4). Moreover, if tadpole size strongly predicted adult reproductive success, then above a certain size difference the smaller tadpole should let its larger relative win without fighting (middle row of panels in Fig. 4).

## Conclusions

In this study, we attempted to better understand the roles of physical attributes and relatedness in predicting aggressive behavior under stable conditions. We found that large siblings were significantly less aggressive than large non-siblings towards their smaller counterpart, demonstrating evidence for kin discrimination in *D. tinctorius*. These findings are complicated by latency to aggression, which showed different trends based on dyad relatedness, although the evolutionary explanation for this pattern remains unclear. We contribute to the growing body of literature demonstrating aggressive behavior (that may escalate to cannibalism) is independent of environmental stress, and conclude that (1) size and relatedness interact in predicting aggressive behaviors, and (2) the ability to discriminate kin does not guarantee kin bias. These findings set the stage for studies to consider cannibalistic behavior in more complex ways, and to better understand the value and purpose of kin discrimination in cannibals.

## Supporting information

Supplementary Table 1. AIC model selection for (A) post-acclimation and (B) experimental activity trials.

Supplementary Table 2. AIC model selection for total aggression trials.

Supplementary Figure 1. Difference in aggression between tadpole dyads.

Supplementary Figure 2. Illustration of Parental mating network for experimental tadpoles.

## Acknowledgements

We thank Teemu Tuomaala for taking care of tadpoles and the lab population; a big shout-out to Aislyn Keyes for advice on coding the mating network which helped generate random pairs to keep our frogs happy and healthy. Enormous thank you to Benjamin Leary for his amazing illustration depicting tadpole aggression. This study was funded by the Academy of Finland (Academy Research Fellowship No.21000042021 to BR).

## Statement of Authorship

BR conceived the study, with input from CF and JV in the design of the assays; CF performed the study; LF developed the mathematical model; CF analyzed the data; and CF wrote the first draft with input from BR. All authors critically commented on previous drafts and approved the final version of the manuscript.

