## Supplementary Table 1. AIC model selection for (A) post-acclimation and (B) experimental activity trials. for "Size-dependent tradeoffs in aggressive behavior towards kin"

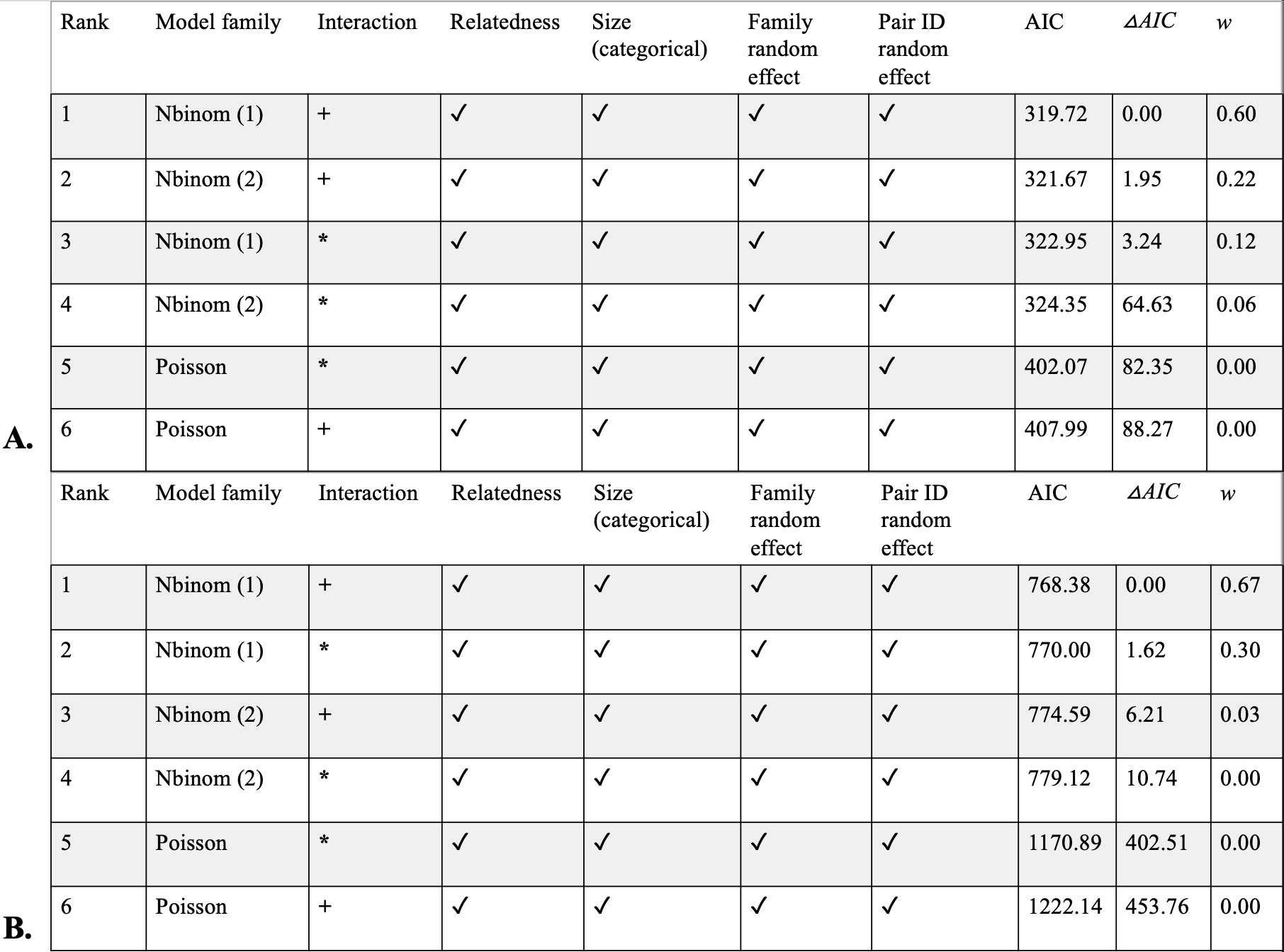

Supp. Table 1. AIC model selection for (A) post-acclimation and (B) experimental activity trials where models with the lowest AIC were selected. Checkmarks indicate predictors included in the model. All models had the same response variable which was activity count. “+” indicaticates additive predictors while “*” indicates interactive terms. AIC is the Akaike Information Criteria and ΔAIC indicates the difference in model support, *w* indicates model probabilities indicating the level of support. All models were coded using a general linear mixed model framework using a template model builder (glmmTMB). Model families: Nbinom1 = Negative binomial distribution: linear parameterization, Nbinom2 = Negative binomial distribution: quadratic parameterization, Poisson = Poisson distribution; both families were considered because event occurrences could be explained by both distributions. Differences in experimental activity (B) were offset by trial durations.
