## Supplementary Table 2. AIC model selection for total aggression trials. for "Size-dependent tradeoffs in aggressive behavior towards kin"

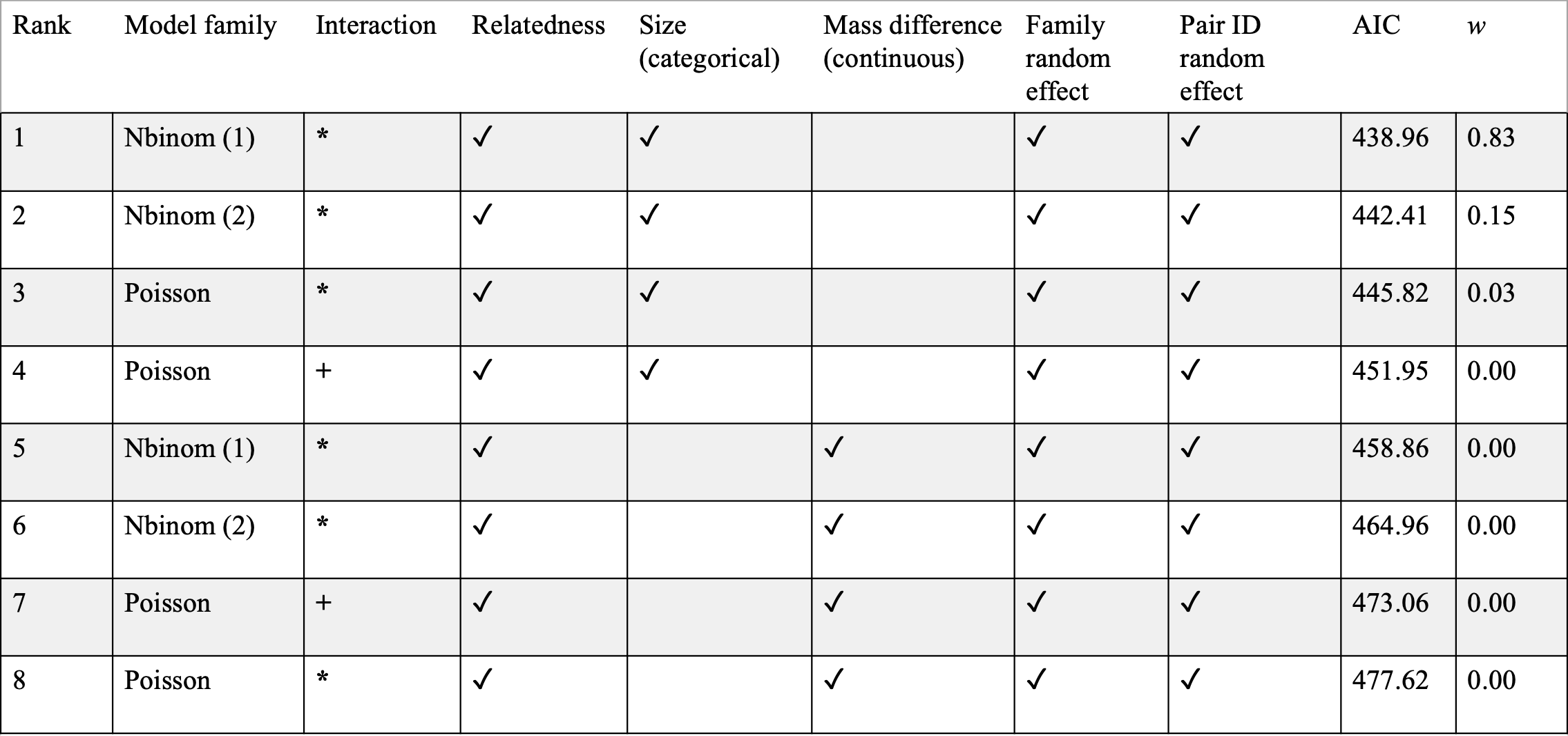

Supp. Table 2. AIC model selection for total aggression trials. Checkmarks indicate predictors included in the model. All models had the same response variable of total aggression, which was the sum of biting and count events during each trial. “+” indicaticates additive predictors while “*” indicated interactive terms. AIC is Akaike Information Criteria, *w* indicates model probabilities indicating the level of support. All models were coded using a general linear mixed model framework using a template model builder (glmmTMB). Model families: Nbinom1 =Negative binomial distribution: linear parameterization, nbinom2 =Negative binomial distribution: quadratic parameterization, Poisson = Poisson distribution; both families were considered because event occurrences could be explained by both distributions. Differences in experimental time were offset by trial durations. Size was a categorical variable where size was relative between dyads (large/small) and mass_diff was the mass difference between the large and small tadpole.
