## Supplementary Figure 1. Difference in aggression between tadpole dyads. for "Size-dependent tradeoffs in aggressive behavior towards kin"

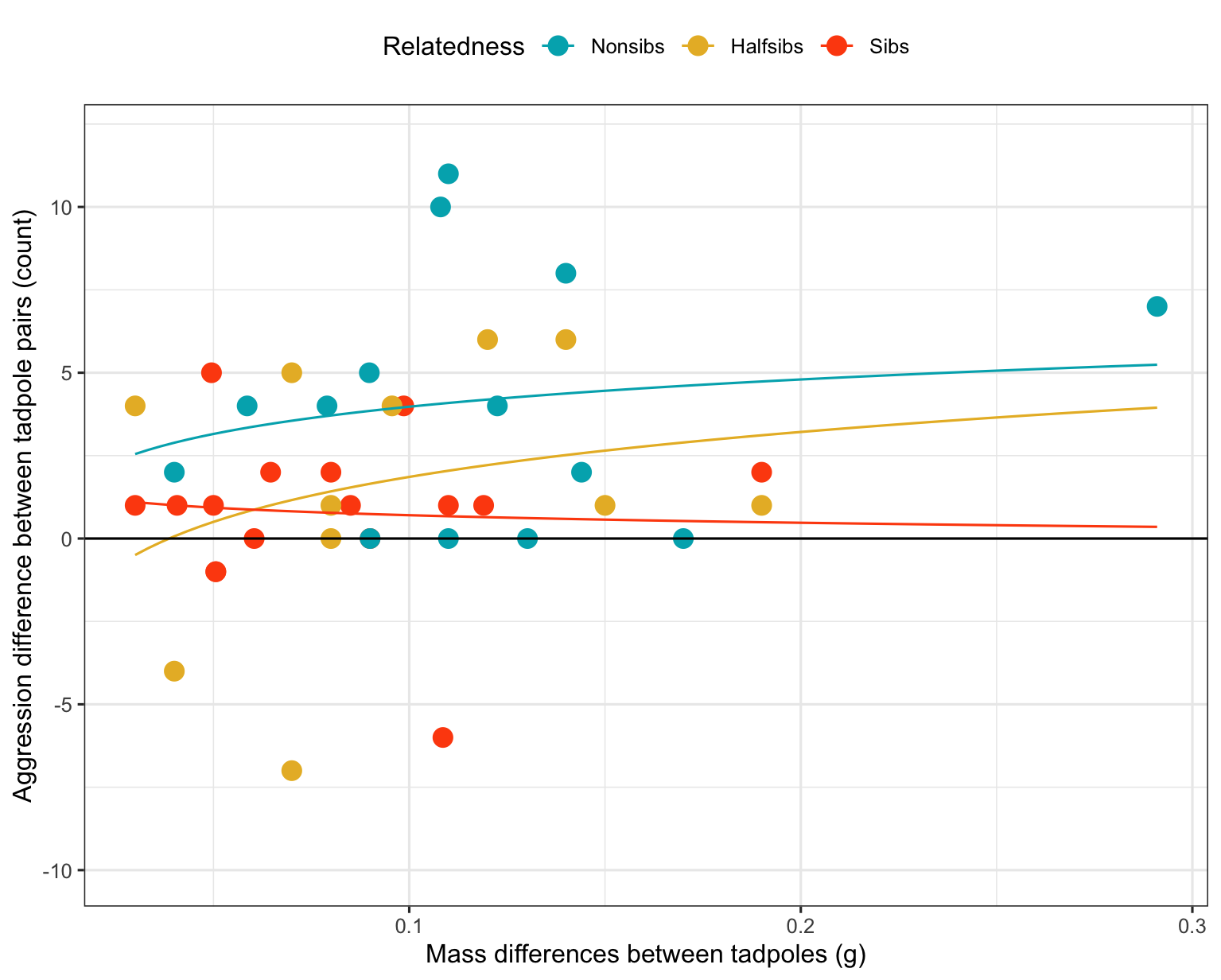

Supp. Fig 1. Difference in aggression between tadpole dyads. All counts are relative to the large tadpole, numbers below zero indicate higher levels of aggression by the small tadpole. GLM smoother fit with a y ~ log(x) formula.
