## Supplementary Figure 2. Illustration of Parental mating network for experimental tadpoles. for "Size-dependent tradeoffs in aggressive behavior towards kin"

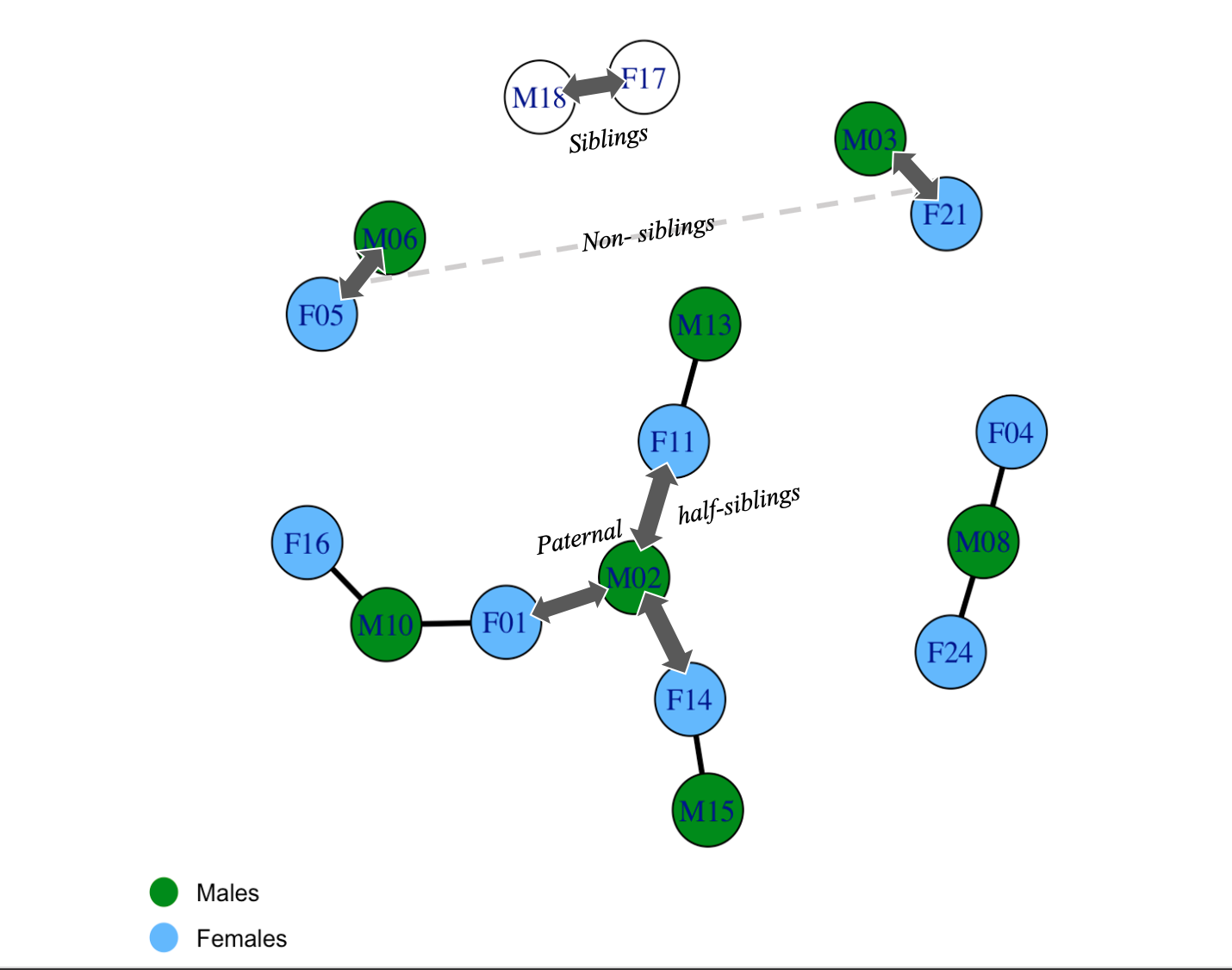

Supp Fig. 2. Illustration of Parental mating network for experimental tadpoles. Males are marked in green circles and females are marked in blue circles. Solid lines indicate successful mating between pairs. Only paternal half-siblings were tested together to avoid possible confounding parental effects. Double headed arrows indicate examples of each of the relatedness classes.
